# Roles and mechanisms of leptomeningeal collaterals in human distal MCA territory ischemia

**DOI:** 10.1101/2024.05.13.594043

**Authors:** Xiang Zou, Shuhao Mei, Zehan Wu, Kun Song, Yuhao Xu, Jie Hu, Liang Chen, Ying Mao

## Abstract

**Background:** Leptomeningeal collaterals (LMCs) provided hemodynamic support and reperfusion during acute ischemic stroke (AIS). Although regulation details have been deeply acquired from animal models, human cortex may share limited hemodynamic patterns due to larger physical scale that needs further investigation.

**Methods:** We performed ‘sequential hierarchical blocking’ during awake craniotomy on fourteen human subjects to mimic AIS in middle cerebral artery (MCA) M_4_ segment territory. Widefield microscope was applied to reveal microcirculation of LMCs. Relative flow rate (RFR) was measured to evaluate the impact of LMCs hierarchy on microcirculation maintenance. Single-cell sequencing and spatial transcriptomics were further performed to analyze the mechanism of different LMCs recruitment abilities.

**Results:** LMCs RFR decreased to 30.75% immediately (95% CI, 6.64 - 54.86%, P < 0.001), and about 80.25% RFR was finally recovered within 5 minutes (95% CI, 55.2 - 105.3%, P = 0.0034) during M_4_ occlusion. For M_5_ and pial branches occlusion, RFR decreased to 55.78% with no further recovery (95% CI, 22.50-89.05%, P = 0.014), and 4 in 11 subjects had no valid microcirculation. For subjects with better recruitment ability, the proportion of arteriole is much higher in leptomeninx (76 - 79% vs. 36 - 56%). There were more significant interaction pairs associated with key signaling pathways between arteriole and neuron/astrocyte (TGF-beta signaling pathway, ECM-receptor interaction)

**Conclusions:** Abundant LMCs supported hemodynamic stability during AIS in human subjects. The proportion of arteriole is much higher in leptomeninx for subjects with better recruitment ability. The Active neuron/astrocyte - arteriole interaction may improve LMCs recruitment.

**GRAPH ABSTRACT:** 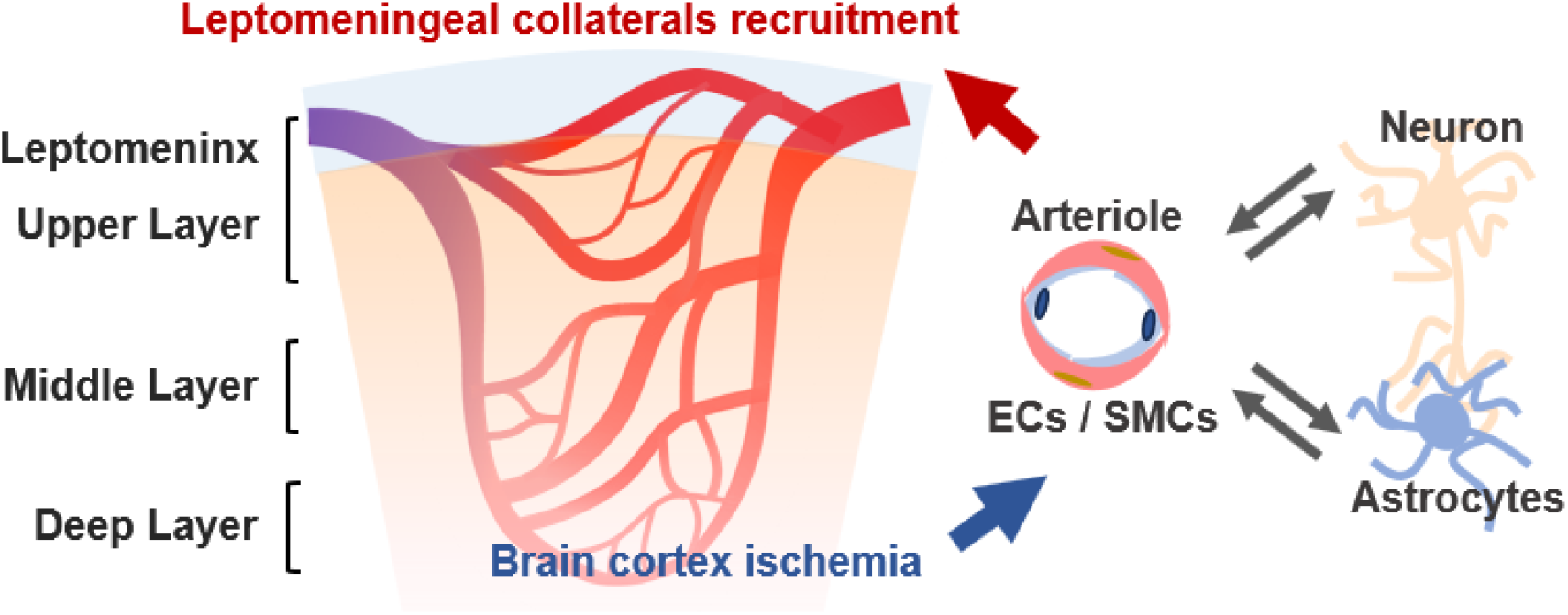

## INTRODUCTION

Incomplete reperfusion of the microvasculature is about eighty percent of acute ischemic stroke (AIS) patients, even after successful recanalization of the occluded vessel^1^. It has been verified in rodent models that abundant leptomeningeal collaterals (LMCs) provided hemodynamic support during AIS and reperfusion, acted as key components regulating reperfusion and preventing futile recanalization^2^. Although the regulation details in microcirculation have been deeply acquired from animal models, human cortex may share limited hemodynamic patterns due to the thicker layers, more collateral networks and larger physical scale. We recently conducted an *in vivo* study on human subjects to investigated the roles and mechanisms of leptomeningeal collaterals in distal middle cerebral artery (MCA) territory ischemia.

## MATERIALS AND METHODS

### Study Design

This study was approved by the Ethics Committee of Huashan hospital, Fudan university (KY2022-591, KY2021-918) and was conducted in accordance with the Declaration of Helsinki (1964) and later amendments, in department of neurosurgery, Huashan hospital, from May, 2023 to Dec, 2023. We performed a procedure called ‘sequential hierarchical blocking’ during extend resection or lobectomy to mimic AIS in MCA M_4_ segment territory, which is defined as step-by-step occluding M_4_ - M_5_ - pial artery (PA), shown in Figure 1A. We used a handheld miniaturized epi-fluorescence widefield microscope (MEW-M) to reveal more details about the microcirculation of LMCs, relative flow rate (RFR) was calculated by real-time flow rate dividing initial flow rate, blood flow rate was measured based on inbuilt software. High-density electrocorticography (HD-ECoG) was applied to record *in situ* local field potential (LFP). In four subjects with different LMCs recruitment abilities, single-cell sequencing and spatial transcriptomics were further performed to analyze the mechanism of LMCs recruitment abilities.

**Figure 1.**
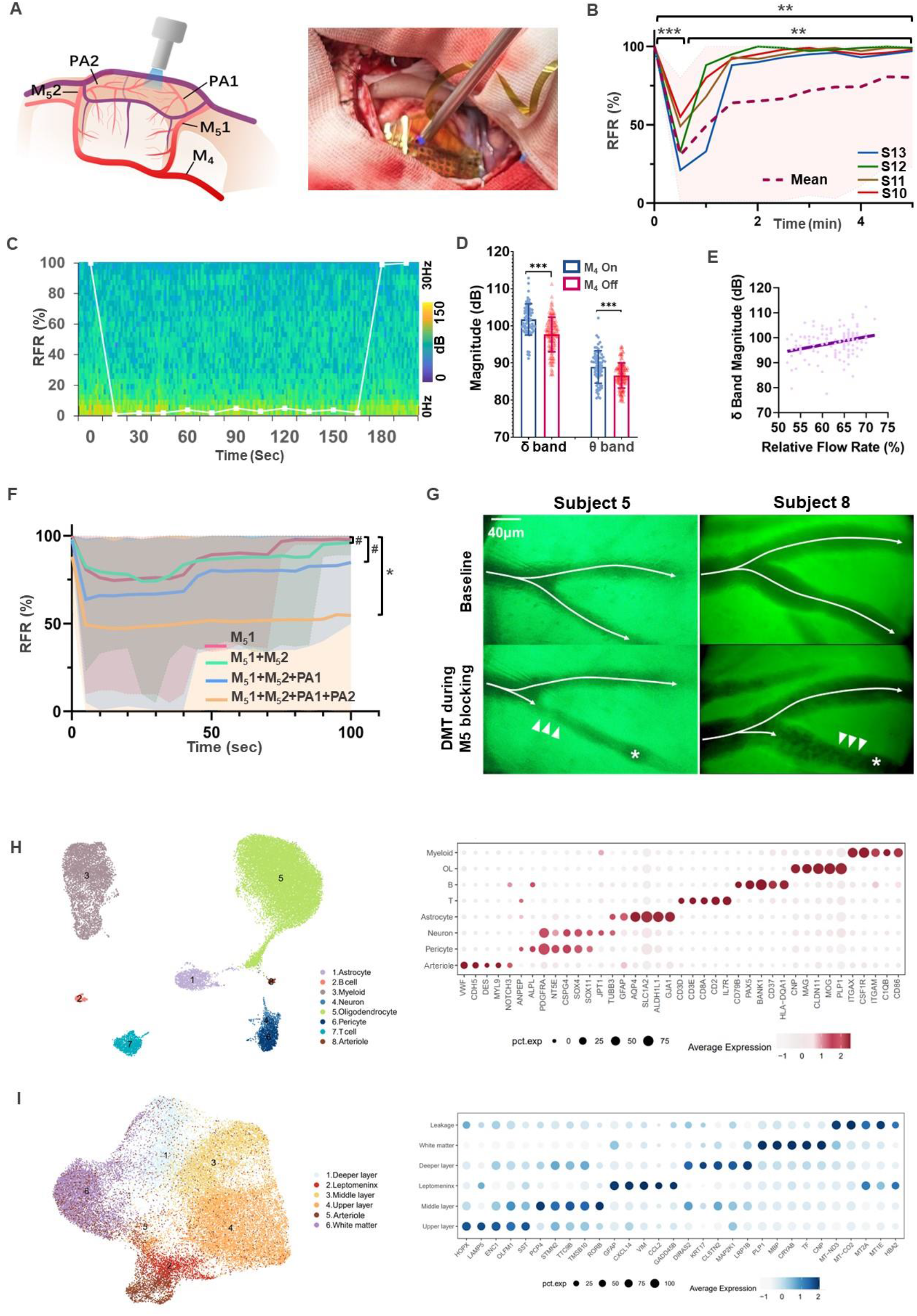
Roles of leptomeningeal collaterals in human distal MCA territory ischemia. (A) Illustration of hierarchical blocking of M4 territory with intraoperative demonstration. (B) RFR after M4 blocking by mean (purple dot) with range (pink area). Patients 10-13’s RFR were highlighted. n = 9, **P < 0.01, ***P < 0.001, Mann-Whitney U test. (C) Correlations of in situ ECoG with RFR after M4 blocking. White broken line: RFR. (D) Correlation analysis of RFR and in situ ECoG band, presented by mean ±SD. ***P < 0.001, unpaired t test. (E) Linear regression analysis between RFR and PSD for δ band. (F) RFR after hierarchical sequence blocking presented as mean (line) with range (colored area). n = 11, *P < 0.05, # P > 0.05, Mann-Whitney U test. (G) Downstream microvascular thrombosis for subject 5 and 8. Arrows: flow direction; triangle: vasospasm; asterisk: microemboli. Scale bar: 40 μm. (H) UMAP plots of cortical cell types of subjects 10 to 13 (left). Bubble plots of the top 5 differentially expressed genes (right). (I) UMAP plots of various cortical layers of subject 10 to 13, as well as the arteriole plots into 5 different cortical layers (left). Bubble plots of the top 5 differentially expressed genes (right).

### Participants

This case series study was approved by the Ethics Committee of Huashan hospital, Fudan university (KY2022-591, KY2021-918) and was conducted in accordance with the Declaration of Helsinki (1964) and later amendments, in department of neurosurgery from May, 2023 to Dec, 2023. Inclusion criteria were patients with low grade glioma or epilepsy, frontal/temporal lobectomy or extended resection will be performed according the preoperative plan under awake craniotomy. Thus, we will have normal gyrus to study before removal without hemodynamic interference by anesthetic agent. Exclusion criteria were: tissue edema or abnormal angioarchitecture in the target gyrus confirmed by magnetic resonance imaging (MRI).

### Intraoperative devices

MEW-M used in this study was produced by Dendrite Precision Instruments Inc.t, and HD-ECoG was produced by NeuroXess (Shanghai Brain Tiger Technology Co., Ltd.). Both of them did not participate in the study and interfered the result. Leptomeninges for observation was stained with 0.2% fluorescein sodium for 30 seconds, rinsed with saline.

### Sequential hierarchical blocking procedures

Sequential hierarchical blocking strategy is defined as step-by-step occluding M_4_-M_5_-PA, shown in Figure. 1A, which is: 1) the target gyrus in the territory was chosen following craniotomy, M_4_ was identified and dissected as a standby; 2) the distal M_5_ and pial arteries were identified; 3) high-density electrocorticography (HD-ECoG) was applied to record local field potential (LFP) for power spectral density (PSD) analysis under appropriate conditions^3,4^; 4) Pial arteriole (PA) under 40 μm was chosen by MEW-M system for microcirculation evaluation. Initial flow rate was recorded without any interference; 5) M_4_ was temporally occluded by an aneurysm clip for 5 minutes, then remove the clip. During HD-ECoG recording, M_4_ clipping and repetency was performed several times; 6) M_5_1-M_5_2-PA1-PA2 occlusion was performed step-by-step using aneurysm clip or low power electrocoagulation, in order to mimic the ischemic stroke progress. If M_4_ could not be dissected, the procedure would begin at this step; 7) for each artery occlusion, local microcirculation was observed about 3 minutes till the blood flow became stable; 8) blood flow rate was measured based on inbuilt software.

### Comparison and Quantification of Stereo-seq and 10X Data

We performed automated image quality control using ImageStudio and processed the raw data with the SAW6.1 workflow (STOmics/SAW [github.com]), using the hg38 reference genome from the Ensembl database (fa) and the accompanying GTF annotation file, built with STAR^5^. The resulting files were input into Stereopy (Usage Principles - Stereopy) and converted to formats compatible with downstream requirements. For 10X data, upstream alignment was conducted using Cell Ranger 7.0 (ceranger count function) based on the Human reference (GRCh38) - 2020-A, with default alignment parameters (Running Cell Ranger count - Official 10x Genomics Support).

### Quality Control

To maximize the utility of our Stereo-seq data, we experimented with analysis from bin50 (50 × 50 DNB bins/spot, equivalent to 25 μm) to bin200, ultimately selecting bin150 for analysis to optimally balance data quality and accuracy. Subsequently, we used Seurat V5.0.3 to retain cells with a mitochondrial gene proportion less than 20%, and nFeature between 500 and 10000^6^. For the 10X sequencing results, we initially employed default parameters of DoubletFinder to remove doublets for each sample separately, retaining cells with mitochondrial proportions less than 20% and nFeature between 500 and 5000^7^.

### Cell Type Discrimination

We utilized the default settings of STAGATE to differentiate various cortical layers of spatial data, integrating slice spatial information for cortical segmentation. Next, we employed the FindAllMarkers function of Seurat to identify differentially expressed genes (DEGs) among cortical layers, and created bubble plots of the top 5 DEGs using custom R functions. For scRNA-seq data, we integrated classic markers from CellMarkersand multiple references to distinguish neural, immune, and vascular cells, visualizing these markers in bubble plots^8-12^.

### Cell Type Differentiation

We utilized the default settings of STAGATE to differentiate various cortical layers of spatial data, integrating slice spatial information for cortical segmentation. Subsequently, we employed Seurat’s FindAllMarkers function to identify differential genes between these cortical areas. Custom R functions were used to generate bubble charts for the top 5 differentially expressed genes (DEGs). For scRNA-seq data, batch integration was performed using Harmony, followed by principal component analysis (PCA) and UMAP dimensionality reduction using Seurat’s default parameters. Multiple resolutions in the FindClusters function, classic markers from CellMarkers, and additional markers derived from several references were utilized to identify neuronal, immune, and vascular cells. Bubble charts were created using those markers^8-12^.

### Mapping scRNA-seq Data to Spatial Data

To explore the distribution of annotated vascular cells across brain cortical layers, we employed Seurat’s MapQuery function to map vascular cells onto spatial data. We further utilized custom R functions to generate stacked bar charts displaying the distribution proportion of arteriole across four samples in each cortical layer.

### Cell Communication Analysis

We conducted communication analysis on scRNA-seq data using CellChat^13^, employing the entire CellChatDB.human database. For visualizing with netVisual_chord_gene, we set additional parameters with a threshold of 0.01, while otherwise using default settings.

### Statistics

Relative flow rate (RFR) was calculated by real-time flow rate dividing initial flow rate. Comparison of RFR between two groups was made using nonparametric Mann-Whitney U (unpaired). Calculation of PSD magnitude in different ECoG bands was previously reported^4^. PSD before M4 blocking and repetency was compared by Student’s t test. The association between PSD magnitude and RFR was further verified by linear regression. Statistical analyses were performed using by GraphPad Prism (version 9.0.0; GraphPad Software, LLC).

## Data Availability

Spatial transcriptomics data are available for qualified researchers upon request to the corresponding authors.

## RESULTS

### Roles of leptomeningeal collaterals in human distal MCA territory ischemia

Fourteen patients with low grade glioma or epilepsy under awake craniotomy were included (Table S1). During the procedure, M_4_ was firstly occluded by an aneurysm clip for 5 minutes. We found that relative flow rate (RFR, real-time flow rate divides by baseline) of LMCs decreased to 30.75% immediately (95% CI, 6.64 - 54.86%, *P* < 0.001), and about 80.25% RFR was finally recovered within 5 minutes (95% CI, 55.2 - 105.3%, *P* = 0.0034), but not completely compared to beginning (*P* = 0.0014, Figure 1B). Among these subjects, the immediate drop of RFR can range from 20 - 99%, and the recovery of RFR can range from 23 - 100%, indicating different LMCs recruitment abilities.

During M_4_ clipping and repetency, we found *in situ* PSD magnitude of δ (0 - 3 Hz) and θ bands (4 - 7 Hz) were closely related to LMCs’ RFR (101.7 ±4.2 vs. 97.7 ±4.7 dB, *P* < 0.001; 88.9 ±4.4 vs. 86.6 ±3.4 dB, *P* < 0.001, Figure 1C), while δ band was positively correlated with LMCs’ RFR (R^2^ = 0.09, *P* < 0.001, Figure 1D), representing the neural activity related to LMCs recruitment.

To mimic AIS progress, M_5_1-M_5_2-PA1-PA2 occlusion was then performed. We found that M_5_ and pial branches were more critical for microcirculation maintenance: mean RFR decreased to 55.78% with no further recovery (95% CI, 22.50-89.05%, *P* = 0.014), and 4 in 11 subjects had no valid microcirculation when finished the procedure (Figure 1E). Notably, we found downstream microvascular thrombosis (DMT) could appear and block arteriole of LMCs completely during proximal occlusion in two cases (18.1%), even if the homologous branch still maintained its 20 - 70% RFR (Figure 1F, online supplemental video S1 and S2).

### Mechanism of different LMCs recruitment abilities analyzed by spatial transcriptomics

We further hypothesized that LMCs recruitment ability might rely on arteriole distribution in the cortex. Main cell types were defined in cortex from subject 10 to 13 (Figure 1H)^8-12^, including arteriole marked by endothelial cells (*VWF, CDH5, DES*) with smooth muscle cells (*MYL9, NOTCH3*). Then cortex was layered from leptomeninx to white matter, and the distribution of arteriole was merged into different layers (Figure 1I). LMCs recruitment abilities were analysis in subject 10 to 13, which were shown better in subject 10 and 11 (more that 50% RFR maintenance after the procedures), while worse in subject 12 and 13 (no recruitment after the procedures, Figure 2A). For subjects with better recruitment ability, the proportion of arteriole is much higher in leptomeninx (76 - 79% vs. 36 - 56%, Figure 2B-C), while the proportion of capillary has no such trend (mostly in the grey matter for all subjects, Figure 2D). Moreover, ligand-receptor interaction pairs between arteriole and surrounding cells clusters including neuron and astrocyte were further analyzed^13^. In subject with better LMCs recruitment, there were greater strength of ligand - receptor pairs between neuron/astrocyte – arteriole, and more significant (p < 0.05) interaction pairs associated with key signaling pathways between arteriole and neuron/astrocyte (TGF-beta signaling pathway, ECM-receptor interaction, Figure 2E). In contrast, such interactions in subjects with worse LMCs recruitment abilities were weaker (Figure 2F). Details were available in Supplemental dataset 1.

**Figure 2.**
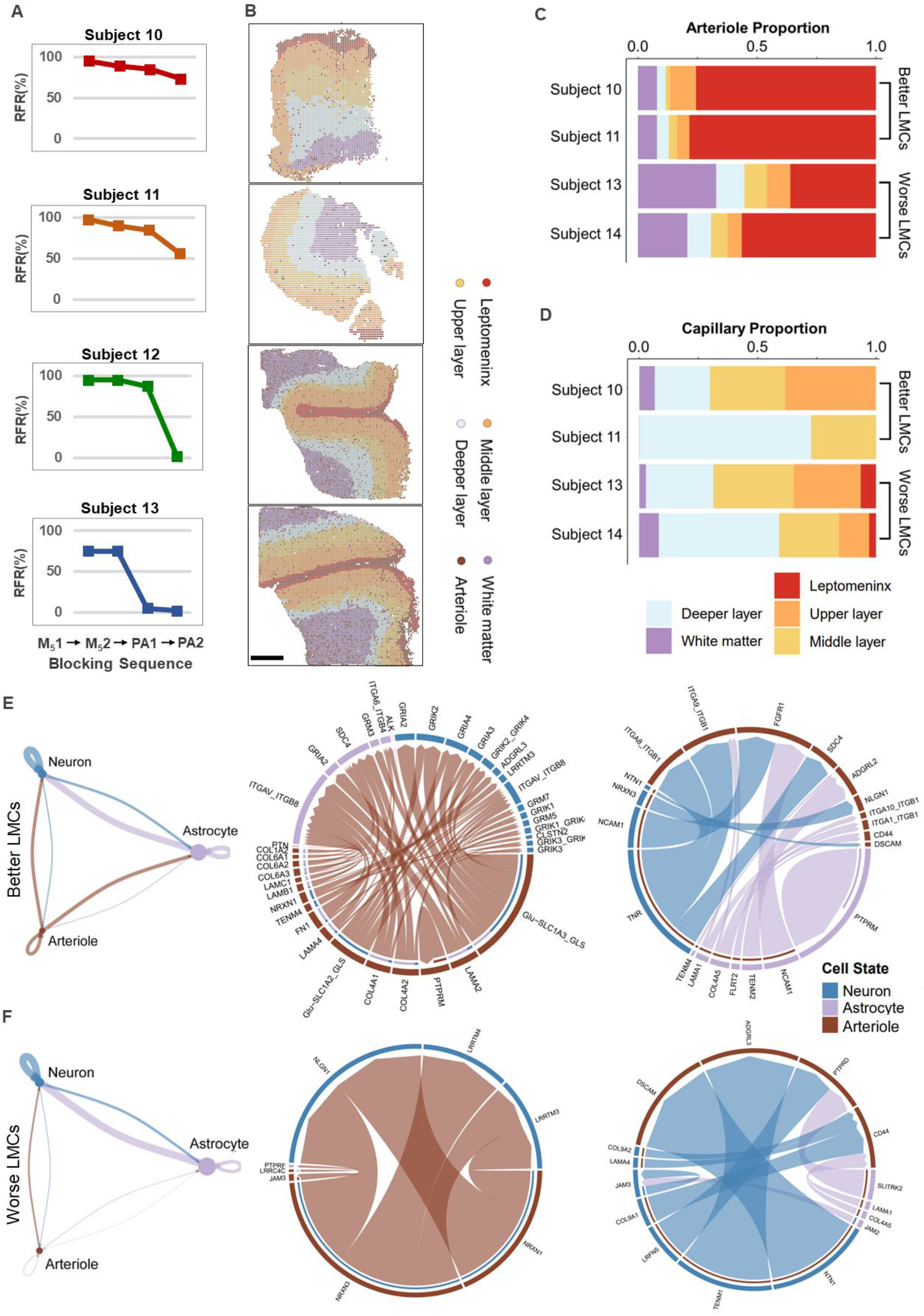
Mechanism of different LMCs recruitment abilities analyzed by spatial transcriptomics. (A) LMCs recruitment abilities of subject 10 to 13 evaluated by M51-M52-PA1-PA2 blocking. (B) Distributions of arteriole across brain cortical layers of subject 10 to 13. Bins/dots are colored by their annotations from Figure 1I. Scale bar: 2 mm. (C) Bar charts displaying the distribution proportions of arteriole across four samples in each cortical layer. (D) Bar charts displaying the distribution proportions of capillary across four samples in each cortical layer. (E) The chord diagram shows the ligand-receptor interaction pairs between arteriole and surrounding cells clusters including neuron and astrocyte, in subject 10 and 11 and in subject 12 and 13 (F). The thickness of the line represents the strength of ligand-receptor pairs. The circos diagrams showing the significant (p < 0.05) interaction pairs associated with key signaling pathways from arteriole to neuron and astrocyte (middle) and from neuron and astrocyte to arteriole (right).

## DISCUSSION

Recent clinical trials held different opinions on thrombolysis combining with thrombectomy^14,15^, which made us rethink the different hierarchical roles in microcirculation maintenance. We provide human *in vivo* evidence that abundant LMCs supported hemodynamic stability during AIS in microcirculation level. The above findings indicated the hierarchical roles along LMCs tree: proximal LMCs arteries are related to the volume of microcirculation, while distal arterioles determine microcirculation existence. As DMT may also play a key role in futile recanalization, we suggest shorter door-to-needle time of thrombolysis could improve prognosis^16^. Those findings are also in agreement with recent clinical trials that suggest thrombolysis or strengthening anti-platelet treatment, either in large or medium-small sized vessel occlusion to improve prognosis^14,15^. Limited by awake craniotomy procedures, only frontal/temporal M_4_ territory was studied, which may not exactly represent the situation in other brain areas. Further studies including large sample size and reperfusion procedures may allow to verify more basic knowledge in human AIS.

In summary, this study demonstrates the arteriole density in leptomeninx and upper cortex layer is more important to brain microcirculation maintenance than other cortex layers. Additionally, we found that interaction of arteriole and surrounding cells improved the effects of LMCs recruitment.

Cerebral vasospasm may be induced by DMT, indicating current vasoactive agent for vasodilation may have less therapeutic effect on incomplete reperfusion without thrombolysis. Future therapeutic strategy for AIS should pay more attention on neuron/astrocyte - arteriole interactions that may improve LMCs recruitment. As LMCs also maintain the brain microcirculation during fluctuation of blood pressure, our findings will also inspire systemic treatment strategy in cardiovascular disease.

## ACKNOWLEDGEMENTS

None.

## SOURCES OF FUNDING

This study was supported by National Natural Science Foundation of China (81970418 from Dr. Zou, 82272116 from Prof. Chen); Science and Technology Innovation Plan of Shanghai Science and Technology Commission (23Y31900300 from Prof. Mao, 21Y21900600 from Prof. Hu)

## DISCLOSURES

The authors declare no competing interests.

## SUPPLEMENTAL MATERIAL

Tables S1

Videos S1–S2

Data Set 1

## Highlights

1. First in vivo observation and challenge of human LMCs.
2. First spatial transcriptomic study for different LMCs recruitment phenotypes.
3. Abundant LMCs supported hemodynamic stability during AIS in human subjects.
4. The proportion of arteriole is much higher in leptomeninx for subjects with better recruitment ability.
5. The Active neuron/astrocyte - arteriole interaction may improve LMCs recruitment.

## Notes

### Competing Interest Statement

The authors have declared no competing interest.

